# Synchrony of mind and body are distinct in mother-child dyads

**DOI:** 10.1101/2021.02.21.432077

**Authors:** Vanessa Reindl, Sam Wass, Victoria Leong, Wolfgang Scharke, Sandra Wistuba, Christina Lisa Wirth, Kerstin Konrad, Christian Gerloff

**Affiliations:** Child Neuropsychology Section, Department of Child and Adolescent Psychiatry, Psychosomatics and Psychotherapy, Medical Faculty, RWTH Aachen University, Germany; JARA-Brain Institute II, Molecular Neuroscience and Neuroimaging, RWTH Aachen & Research Centre Juelich, Germany; Division of Psychology, University of East London, London E16 2RD, United Kingdom; Department of Psychology, University of Cambridge, Cambridge CB2 3EB, United Kingdom; Division of Psychology, Nanyang Technological University, Singapore S639818, Republic of Singapore; Chair of Cognitive and Experimental Psychology, Institute of Psychology, RWTH Aachen University; Chair II of Mathematics, Faculty of Mathematics, Computer Science and Natural Sciences, RWTH Aachen University, Germany

**Keywords:** interpersonal synchrony, hyperscanning, multimodal imaging, functional near-infrared spectroscopy, electrocardiography

## Abstract

Hyperscanning studies have begun to unravel the brain mechanisms underlying social interaction, indicating a functional role for interpersonal neural synchronization (INS), yet the mechanisms that drive INS are poorly understood. While interpersonal synchrony is considered a multimodal phenomenon, it is not clear how different biological and behavioral synchrony markers are related to each other. The current study, thus, addresses whether INS is functionally-distinct from synchrony in other systems – specifically the autonomic nervous system (ANS) and motor behavior. To test this, we used a novel methodological approach, based on concurrent functional near-infrared spectroscopy-electrocardiography, recorded while *N* = 34 mother-child and stranger-child dyads (child mean age 14 years) engaged in cooperative and competitive tasks. Results showed a marked differentiation between neural, ANS and behavioral synchrony. Importantly, only in the neural domain was higher synchrony for mother-child compared to stranger-child dyads observed. Further, ANS and neural synchrony were positively related during competition but not during cooperation. These results suggest that synchrony in different behavioral and biological systems may reflect distinct processes. Mother-child INS may arise due to neural processes related to social affiliation, which go beyond shared arousal and similarities in behavior.

## 1 Introduction

Historically, the mind and the body are considered distinct in Western philosophy. This dualism however does not hold true in modern sciences ^1^. The brain is an interoperable system which is embedded in the human body and influenced by other biological systems. In accordance with this, neuroimaging studies in individual subjects have shown that fluctuations in the autonomic nervous system (ANS) are coupled with changes in brain activity ^2–5^.

While these studies have examined single subjects, humans are social species, who constantly interact with each other. During social interaction people synchronize on many different levels, including their autonomic physiology, behavior, and neural signals ^6–8^. While interpersonal neural synchrony (INS) has been robustly demonstrated in a variety of interactive tasks, the manifold factors which may lead to or affect INS are still poorly understood. Although several studies have provided first important insights (e.g., ^9–14^), these studies often do not consider that synchrony may be established in different behavioral and biological systems. In particular, very little is known about the relationship between INS and synchrony in other biological systems, such as the ANS.

Although no published hyperscanning study that we know of has measured synchrony in neural and ANS domains concurrently, synchrony in either ANS or brain signals has been found in emotional tasks, such as cooperative and competitive games ^15–18^. Thus, while a relationship seems likely, it remains unclear whether synchrony in both systems co-occurs and affects each other.

To investigate this, the present study uses a well-established hyperscanning paradigm ^9,10,13,15,19,20^, in which adult and child either had to cooperate (to synchronize their reaction times to respond as simultaneously as possible to a signal) or to compete (to try to respond faster than their partner to a signal). Participants were 10-18-year-old children and adolescents (all female) who completed the tasks both with their biological mothers (mother-child dyads) and with a previously unacquainted female adult (stranger-child dyads). Previous research shows significant synchrony across the dorsolateral prefrontal and frontopolar cortex when 5-9-year-old children cooperated with their mother, but not in other conditions (mother-child competition, stranger-child cooperation and competition) ^15^. Consistent results have been observed with adults ^13,19^. With older children and adolescents (8-18-year-olds), however, research using the same paradigm additionally identified significant INS during parent-child competition ^20^. One untested possibility is that developmental changes in adolescence may be associated with more emotional arousal and associated ANS synchrony during competition with the parents, potentially leading to increased INS.

To measure neural synchrony, we used functional near-infrared spectroscopy (fNIRS) which allows us to study social interactions in natural, daily life settings, and is relatively robust against motion. Importantly, it not only captures the brain’s local hemodynamic response with a high temporal resolution, but also provides spatial information to locate brain regions which drive INS (e.g., ^21^). To obtain comparable information about the temporal relationships between two person’s ANS signals, metrics with a high temporal resolution are necessary. The interbeat interval (IBI), that is the time between consecutive heart beats, provides an overall index of arousal, reflective of both sympathetic and parasympathetic activity, which can be assessed reliably within short time windows ^8,22^. Thus, in the current study, we extend fNIRS hyperscanning by using concurrent fNIRS - electrocardiography (ECG) recordings, to measure synchrony in the dyad’s brain signals and IBIs simultaneously. We examined INS in the frequency range of 0.08 to 0.5 Hz, which is outside the range of 1 - 3 Hz that may be contaminated by ECG artifacts.

To analyse INS and its relationship to synchrony in other modalities we developed a new analytical framework based on bipartite graph analyses (described in more detail in Gerloff et al. ^23^). Since complex human behavior and cognition is not localized to a single circumscribed brain region but is organized in functional brain networks, INS may be more accurately modelled as the bidirectional links between the brain networks of interacting subjects (see also ^24,25^). These functional networks can be expressed as graphs. While global graph metrics provide a scalar value, which can be easily compared to synchrony measures in other modalities, nodal metrics provide increased topological detail. Because ANS synchrony might impact INS in very specific brain regions while other nodes might be less affected (see also ^26^), to fully understand whether INS is functionally-distinct from synchrony in other systems, an analysis on both levels may be necessary.

Here, we explored whether INS, measured at both global and nodal levels, goes beyond synchrony in the ANS, as measured by the dyad’s IBIs. We also examined the relationship of INS to behavioral synchrony (indexed as the mean of the absolute differences in response times), and we measured how trial-by-trial adaptations in response times, contingent on feedback during the task, related to INS. To this end, we first compared the different biobehavioral synchrony markers (INS, ANS and behavior), and tested whether, for each measure: i) synchrony was higher for mother-child compared to stranger-child dyads, and ii) synchrony was higher for cooperation and / or competition compared to a non-interactive baseline condition, in which the mother-child / stranger-child dyad watched a relaxing video together (Research Question 1). Second, we explored whether INS was predicted either by ANS synchrony and / or by behavioral synchrony (Research Question 2).

## 2 Results

### 2.1 In which systems does interpersonal synchrony occur?

For the first research question, we examined task (baseline vs. cooperation / competition) and partner (mother vs. stranger) differences in i) INS, ii) ANS synchrony and iii) behavioral synchrony. The subsections are organized as follows. First, we compared mother / stranger-child synchrony to the synchrony of shuffled adult-child pairs, who performed the same task independently of each other. Second, we directly compared the experimental conditions. To analyze the data, we used Bayesian Hierarchical Models (BHMs), deriving a precise probability estimate for each parameter rather than relying on frequentist statistics.

#### 2.1.1 INS

Neural synchrony was assessed over the prefrontal cortex using global and nodal inter-brain density (short ‘density’), which are adapted graph-theoretical measures for hyperscanning (Fig. 1). The nodes of the bipartite graph represent the fNIRS channels of adult and child, respectively, and the edges their connections, quantified by the wavelet coherence. To obtain a more robust network, the number of connections was reduced by a block-wise permutation procedure comparing individual graphs with the graphs of shuffled adult-child pairs. Since many fNIRS hyperscanning studies focus on oxy-hemoglobin (HbO) signals ^9,10,13,15,19^, the results for HbO are presented in the main text and then compared to the results for deoxy-hemoglobin (HbR) to validate the findings and reduce the risk of false positives ^27^ (Supplementary Text 2).

**Fig. 1.**
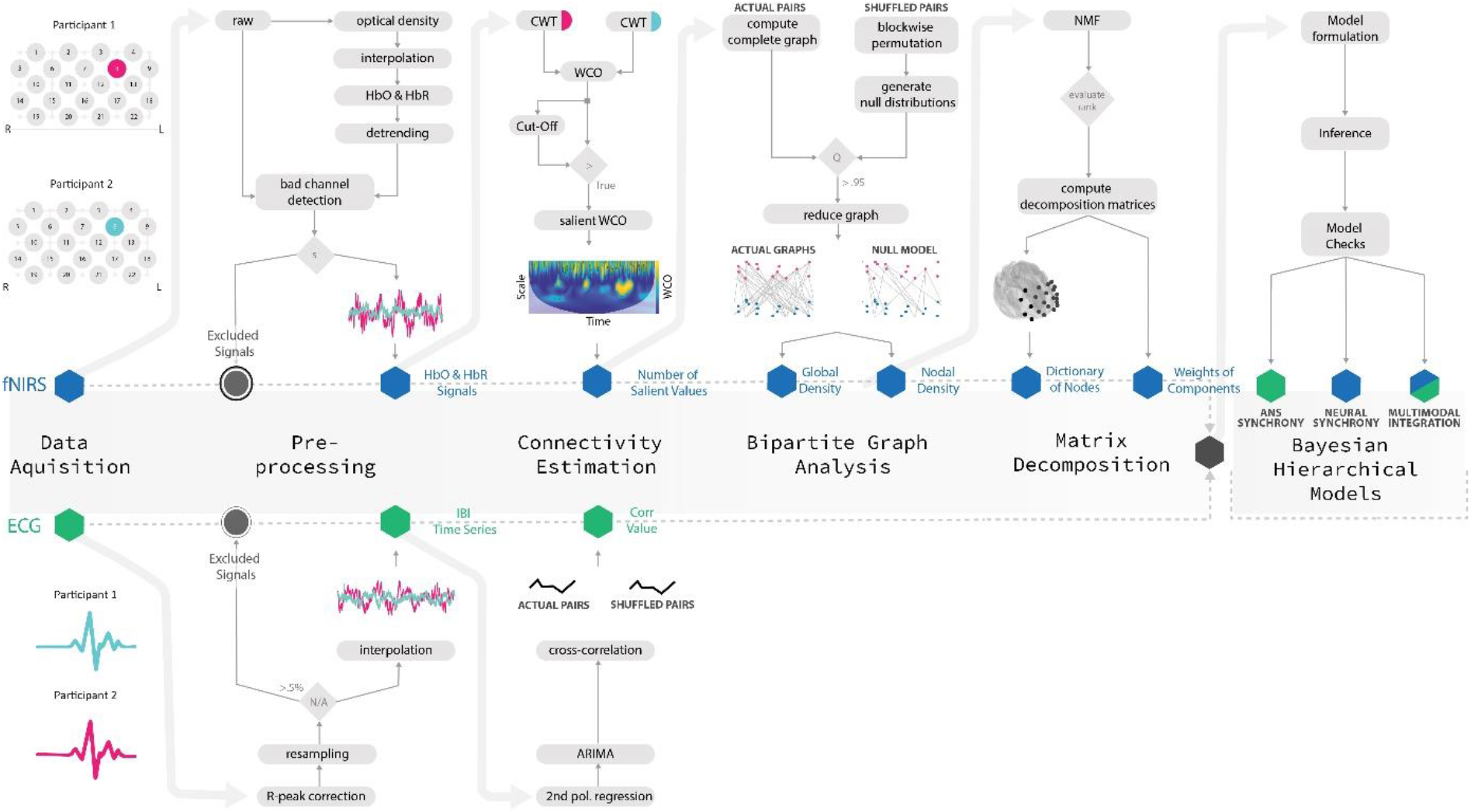
Multimodal data analysis workflow. To examine the relationship between different biobehavioral synchrony measures in a single multivariate generative model, we proposed a symmetric data fusion approach, analyzing synchrony in fNIRS and IBI signals concurrently. Top: After motion artifact correction and detrending of the fNIRS signals, the salient wavelet coherence was calculated as the connectivity estimator. Subsequently, for each dyad and condition, individual bipartite graphs were constructed by defining the salient wavelet coherence as weighted edges connecting different regions (nodes) from adult and child. To avoid spurious connections, the graphs were reduced by a block-wise permutation procedure comparing individual graphs with the graphs of shuffled adult-child pairs. The number of surviving connections between brains was calculated for the network (global density) as well as for each node / fNIRS channel (nodal density). To reduce the dimensionality of the nodal metrics while preserving interpretability, nodal density vectors were encoded via non-negative matrix factorization. Bottom: ANS synchrony was calculated by the cross-correlation of the participant’s IBI time series after R-peak correction and ARIMA modeling. Subsequent analyses were performed using (multivariate) Bayesian hierarchical models.

Since we reduced the bipartite graphs via the 95% quantile of shuffled pairs, consequently, shuffled adult-child pairs had a global density of ~ 5%. To investigate whether global density of actual pairs was actually higher, we estimated the effects of shuffled vs. actual pair per condition within a single BHM. Descriptive results are presented in Supplementary Table 2. For HbO, results showed an increased density only for mother-child competition (posterior mean (μ) = 0.11, 90% credible interval (CI) = [0.02, 0.20]), while no sufficient evidence was found for increased density in the other conditions. However, actual pairs had a lower density in the stranger-child baseline condition (μ = −0.17, CI = [−0.29, −0.05]).

Next, a BHM was calculated for the effects of baseline vs. competition, baseline vs. cooperation and stranger vs. mother as well as their two-way interactions on global density (Fig. 2; Supplementary Table 1). Compelling statistical evidence was found for increased density of mother-child compared to stranger-child dyads (μ = 0.13, CI = [0.04, 0.23]) and of competition compared to baseline (μ = 0.13, CI = [0.02, 0.25]). Furthermore, weaker evidence was found for a positive effect of cooperation compared to baseline (μ = 0.08, CI = [−0.02, 0.18], posterior samples above zero: 91.38 % > 0), while insufficient evidence was observed for an interaction between competition / cooperation and partner.

**Fig. 2.**
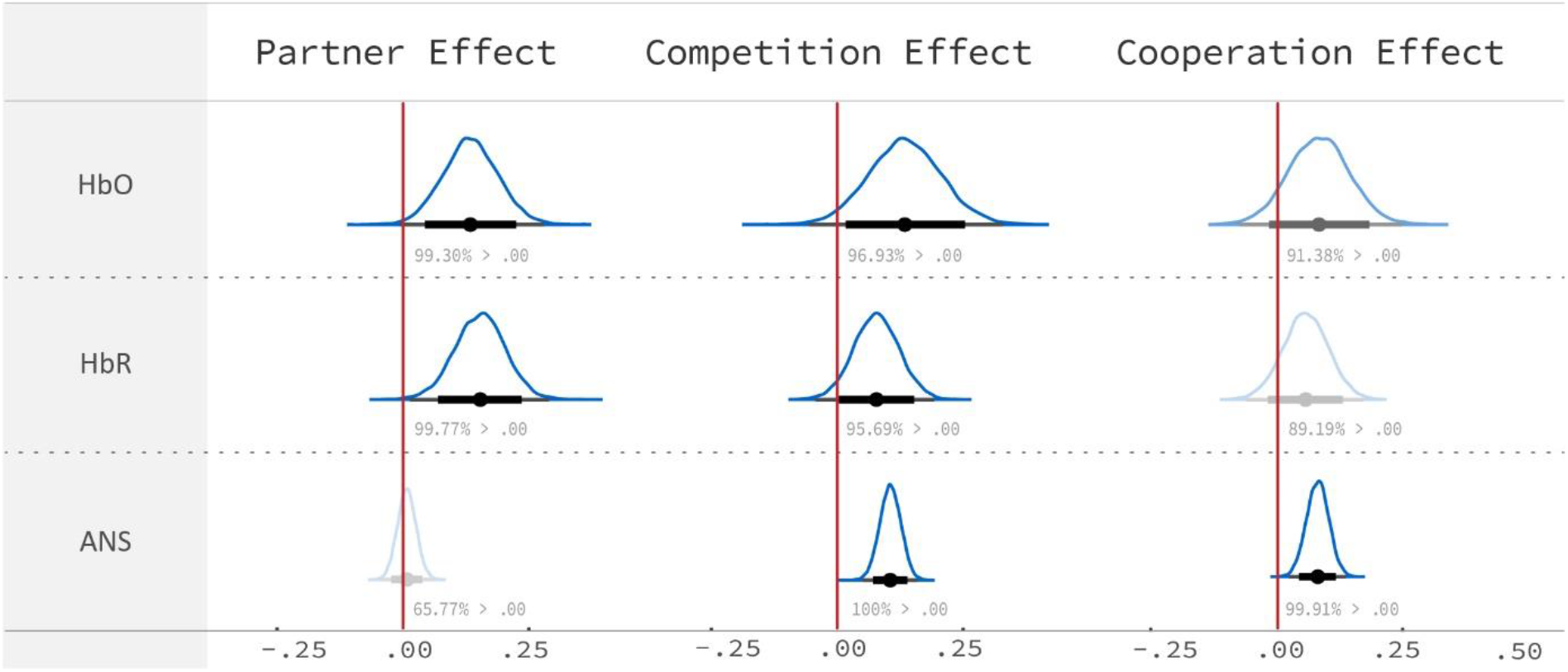
Differences between interpersonal neural and autonomic nervous system (ANS) synchrony as a function of task and partner. To examine the systems in which synchrony occurs, marginal posterior distributions were derived for the effects of stranger vs. mother, baseline vs. competition and baseline vs. cooperation on HbO and HbR global density and ANS synchrony. Forest plots show the 99% and 90% two-sided credible intervals (thin and thick black lines) as well as the posterior mean (black dot). 90% credible intervals which do not cover zero were interpreted as evidence for an effect. For both HbO and HbR, evidence was found for a higher density of mother-child compared to stranger-child dyads and of competition compared to baseline. In contrast, for ANS synchrony, there was no evidence for a partner effect, while strong support was found for both task effects, with increased synchrony for competition and cooperation compared to baseline. Together, these results indicate that synchrony in neural and ANS signals was clearly differentiable.

Since smaller and / or more localized effects may not be detected by global graph metrics, we additionally examined nodal density, having the further advantage of an increased topological detail. The dimensionality of the nodal metrics was reduced via a non-negative matrix factorization (NMF) to four components, each of which contains a collection of nodes with varying contributions. The NMF yields two matrices. The basis matrix can be understood as dictionary to look up the contribution of each node to each component, which is constant across dyads and conditions (Fig. 3). Each node can contribute to several components, just as each brain region can be part of several functional networks. The coefficient matrix encodes the nodal densities of each dyad in the respective condition, with higher values indicating higher INS.

**Fig. 3.**
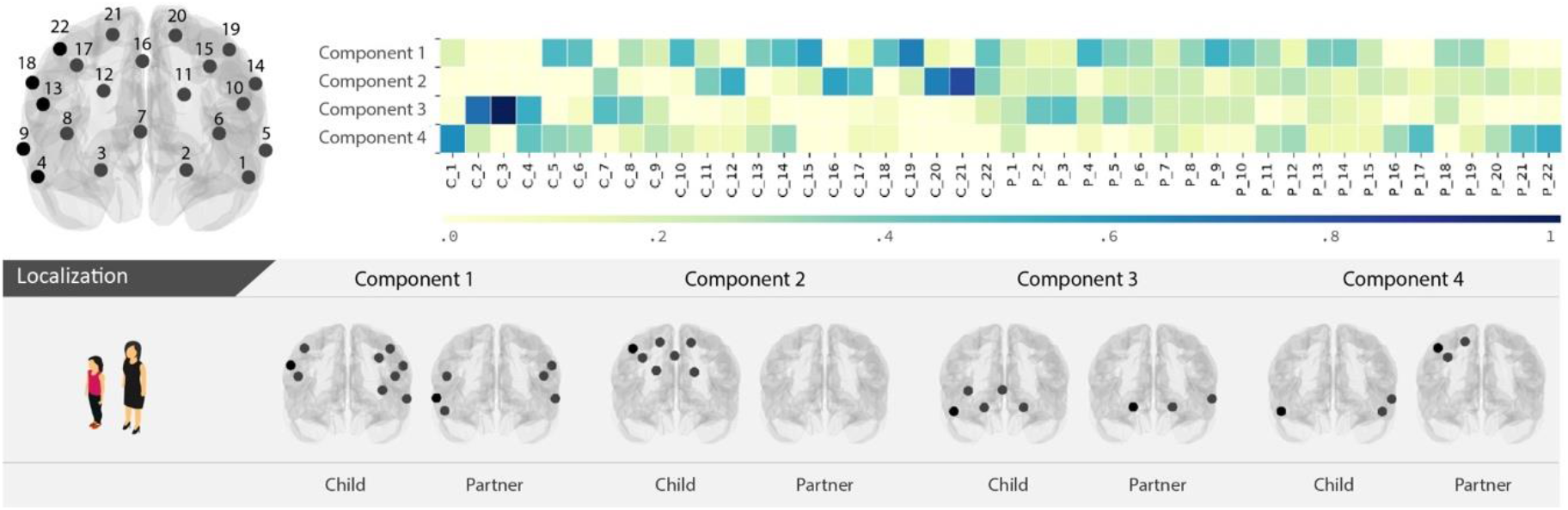
Mapping of NMF components (HbO) to brain regions. Channels and their corresponding MNI coordinates are depicted on the top left. The basis matrix is visualized as a heat map, showing the contribution of each fNIRS channel of child (C) and adult partner (P) (x-axis) to the corresponding component (y-axis). The fNIRS channels of child and adult partner which contribute most to each of the components in terms of their nodal densities, with weights above the 80% quantile (min = 0, max = 1), are depicted on the brains below the heatmap.

Nodal density results validated the global results, showing evidence for a partner, competition and cooperation effect (Fig. 4; Supplementary Table 1). Specifically, we observed higher density for mother-child compared to stranger-child dyads across tasks in component 3 (μ = 0.09, CI = [0.00, 0.18]) and component 4 (μ = 0.15, CI = [0.04, 0.27]) as well as some evidence for an effect in component 1 (μ = 0.08, CI = [−0.02, 0.18], 91.31% > 0). Additionally, in component 2, higher density was found for mother-child compared to stranger-child dyads in the baseline condition (competition x partner interaction: μ = −0.27, 90% CI = [−0.49, −0.05]; cooperation x partner interaction: μ = −0.21, CI = [−0.44, 0.02], 6.32% > 0). Evidence for a competitive task effect was observed in components 1 (μ = 0.16, CI = [0.04, 0.29]) and 4 (μ = 0.13, CI = [0.02, 0.25]) and analogously, evidence for a cooperative task effect was observed in component 1 (μ = 0.12, CI = [0.02, 0.22]) and to a weaker degree in component 4 (μ = 0.09, CI = [−0.02, 0.20], 90.92% > 0). Component 3 and 4 mainly comprise orbitofrontal brain regions of adult and child as well as right superior prefrontal brain regions of the adult, while component 1 and 4 mainly comprise left and right lateralized prefrontal brain regions of adult and child. The brain regions which contribute most to each of the components can be found in Fig. 3.

**Fig. 4.**
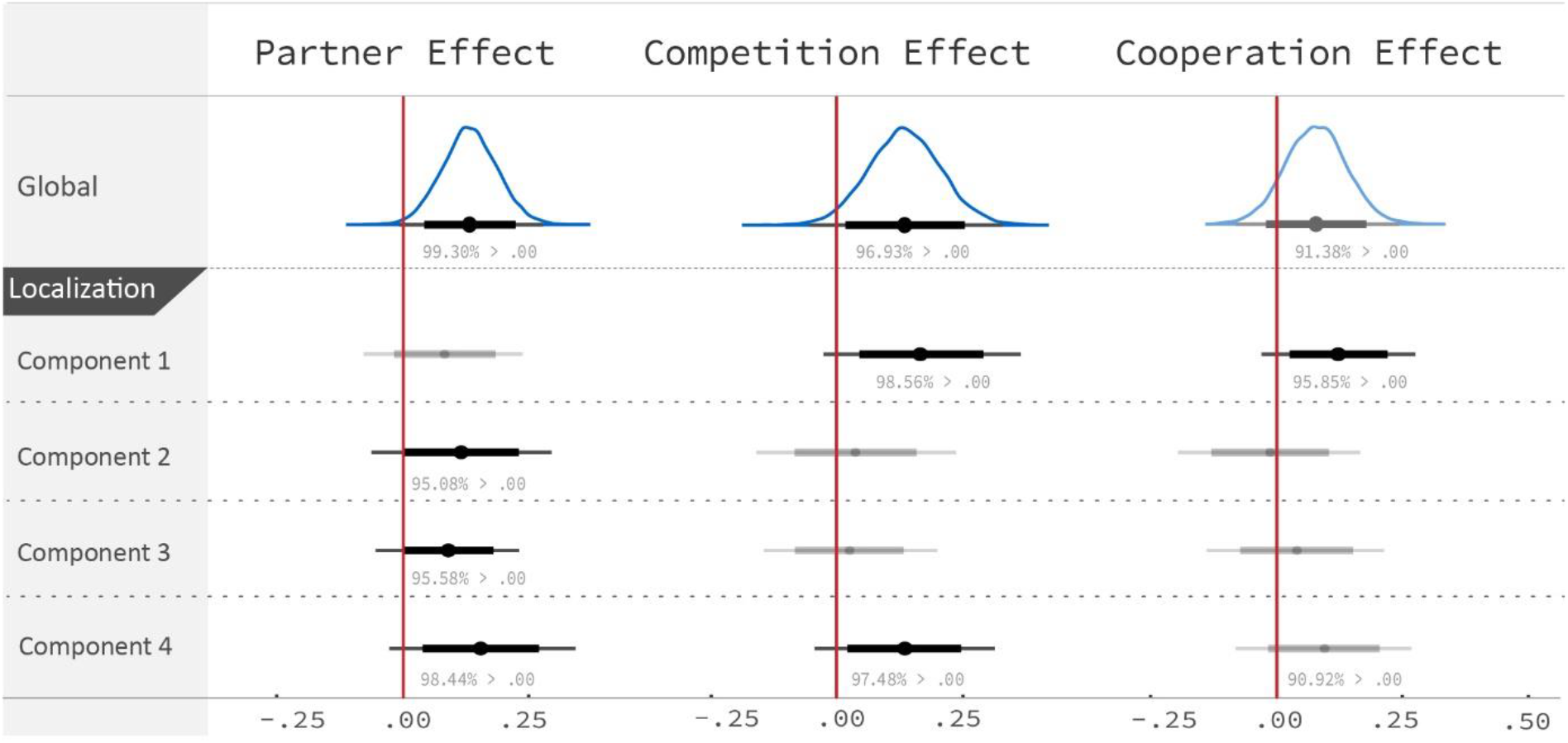
Effects of partner and task on interpersonal neural synchrony measured by global and nodal graph metrics (HbO). In addition to the analyses of global density (Fig. 2), marginal posterior distributions were derived for the effects of stranger vs. mother, baseline vs. competition and baseline vs. cooperation on nodal densities, encoded by the coefficients of the four NMF components. Forest plots show the 99% and 90% two-sided credible intervals (thin and thick black lines) as well as the posterior mean (black dot). Evidence of a partner and competition effect was found both globally and in components 3 and 4 (partner) / components 1 and 4 (competition). Further, a partner effect was found in component 2 however only for the baseline condition. In addition, a cooperative task effect was found in the same components as the competitive task effects, although with weaker evidence. These results show that nodal graph metrics may provide further information on the brain regions which support INS.

Our main neural results (HbO) were further validated by comparing them to the results for HbR, which showed a mostly consistent result pattern (Supplementary Text 2). Again, increased global density was found for mother-child compared to stranger-child dyads and for competition compared to baseline (Fig. 2). Furthermore, in line with the HbO findings, HbR nodal density results confirmed the global findings, indicating increased density for mother-child dyads and for competition. In addition, increased density was found for mother-child cooperation in one component.

Together, these results indicate that INS was increased for mother-child dyads, for competition and for cooperation. Yet, the effects for cooperation were smaller and driven by a subset of nodes as indicated by the NMF results.

#### 2.1.2 ANS synchrony

Partial autocorrelation functions (PACFs) of the ANS data can be found in Supplementary Text 3, Supplementary Figs. 2 and 3. These showed that the data had a strong first-order autoregressive tendency, which motivated our decision to first reduce autocorrelation in the ANS data before calculating the cross-correlation of the IBI time series ^28,29^.

Descriptive results are presented in Supplementary Table 3. When compared to shuffled pairs, increased ANS synchrony was found for mother-child cooperation (μ = 0.08, CI = [0.04, 0.12]), mother-child competition (μ = 0.11, CI = [0.08, 0.14]), stranger-child cooperation (μ = 0.08, CI = [0.04, 0.11]) and stranger-child competition (μ = 0.11, CI = [0.06, 0.15]). However, no increased ANS synchrony was found for mother-child baseline (μ = 0.03, CI = [−0.02, 0.08]) or stranger-child baseline (μ = 0.01, CI = [−0.04, 0.05]).

Directly comparing the conditions, we found very strong evidence for both task effects with higher ANS synchrony for competition (μ = 0.11, CI = [0.07, 0.14]) and cooperation compared to baseline (μ = 0.07, CI = [0.04, 0.11]), although this effect was stronger for competition than for cooperation (μ = 0.03, CI = [0.00, 0.06]). In contrast, no evidence was found for a partner effect and the μ was close to zero (μ = 0.01, CI = [−0.02, 0.04]) (Fig. 2; Supplementary Table 1). Thus, we can conclude with a high certainty that there was increased synchrony for cooperation and competition compared to the non-interactive baseline condition, but no meaningful difference between mother-child and stranger-child dyads.

To ensure that differences between neural and ANS synchrony cannot be attributed to difference in the synchrony estimators, i.e., cross-correlation vs. wavelet coherence, we validated our results by calculating the wavelet coherence on the IBI signals (Supplementary Text 4). In line with the results for the cross-correlation, an increased synchrony was observed for cooperation across dyads. Further, a competition x partner interaction indicated that stranger-child dyads had a higher ANS synchrony for competition compared to baseline, while no task effect was observed for mother-child dyads. In addition, stranger-child dyads had a higher ANS synchrony than mother-child dyads in the competition condition, while no partner effect was observed for cooperation or baseline. These results further demonstrated that increased neural synchrony of mother-child compared to stranger-child dyads was unlikely to be explained by increased ANS synchrony alone.

#### 2.1.3 Behavioral synchrony

Task performance was quantified by first calculating how mean response time differed between conditions (Supplementary Text 5 and Supplementary Table 4). Behavioral synchrony was then measured by calculating the dyad’s mean of the absolute differences in response times (Mean-DRT) during cooperation and competition (Supplementary Table 5). In all conditions, actual pairs were more synchronous than shuffled pairs, although effects were larger for the cooperation conditions (mother-child cooperation: μ = −0.43, CI = [−0.56, −0.31]; stranger-child cooperation: μ = −0.45, CI = [−0.57, −0.33]; mother-child competition: μ = −0.25, CI = [−0.34, −0.16]; stranger-child competition: μ = −0.21, CI = [−0.30, −0.11]). Thus, these findings showed that reaction times of mother / stranger and child were not independent of each other, i.e., Mean-DRTs of actual pairs were smaller than of shuffled pairs.

Directly comparing the conditions, BHM results yielded strong statistical evidence for a task x partner interaction (μ = 0.23, CI = [0.05, 0.41]) (Fig. 2; Supplementary Table 1). Breaking down the interaction, we found that both mother-child and stranger-child dyads were more synchronous during competition than during cooperation (mother-child: μ = −0.32, CI = [−0.48, −0.17]; stranger-child: μ = −0.55, CI = [−0.65, −0.45]). Yet, stranger-child dyads were more synchronous than mother-child dyads during competition (μ = 0.24, CI = [0.11, 0.37]), while no partner differences were found for cooperation (μ = 0.02, CI = [−0.14, 0.17]).

In a supplementary analysis, we examined whether participants adapted their response times after receiving feedback on who had responded more quickly or more slowly (for cooperation this was only the case in trials in which the dyad had lost a point). To this end, differences were calculated between the mean-DRT of the present trial and the mean-DRT of its subsequent trial, whereby larger values indicate a stronger adaptation of the dyad (Supplementary Text 6, Supplementary Table 6). As expected, dyads adapted their RTs more strongly during cooperation than during competition. No associations between these behavioral measures and INS or ANS were observed (Supplementary Tables 7 – 10).

#### 2.1.4 Summary

To summarize, for research question 1, we analyzed whether interpersonal synchrony was observed in multiple systems: in neural signals, ANS, and motor behavior. While results indicated that synchrony was established in all three systems, they also showed that these different synchrony markers were differentially responsive to experimental manipulation. Importantly, only at the brain level mother-child attunement was observed, while no evidence was found for specific attunement in the mother-child dyads’ movements or ANS responses.

### 2.2 Which factors predict interpersonal neural synchrony?

For the second research question, we examined whether task and partner effects on neural synchrony were moderated by ANS and behavioral synchrony. Non-parametric Spearman correlations between the different measures are presented in Supplementary Tables 7 - 12.

To examine the relationship to *ANS synchrony*, BHMs were calculated with the main and interactive effects of baseline vs. competition, baseline vs. cooperation as well as stranger vs. mother with ANS synchrony as predictors and INS as response variable (Fig. 5; Supplementary Table 1). Evidence was found for an interaction of ANS synchrony with baseline vs. competition on global density (μ = 0.91, CI = [0.13, 1.68]), but no sufficient evidence was found for interactions with baseline vs. cooperation and stranger vs. mother. Further analyses of this interaction revealed evidence for an effect of ANS synchrony on global density only for competition (μ = 0.65, CI = [0.21, 1.08]), but not for baseline or cooperation, showing that during competition, higher ANS synchrony predicted increased INS. This should however not be interpreted as a casual or directional effect but rather as an association which could possibly be bi-directional in nature. In line thereof, we also checked the reverse relationship, confirming that increased INS also predicted increased ANS synchrony during competition in our statistical model (Supplementary Text 7).

**Fig. 5.**
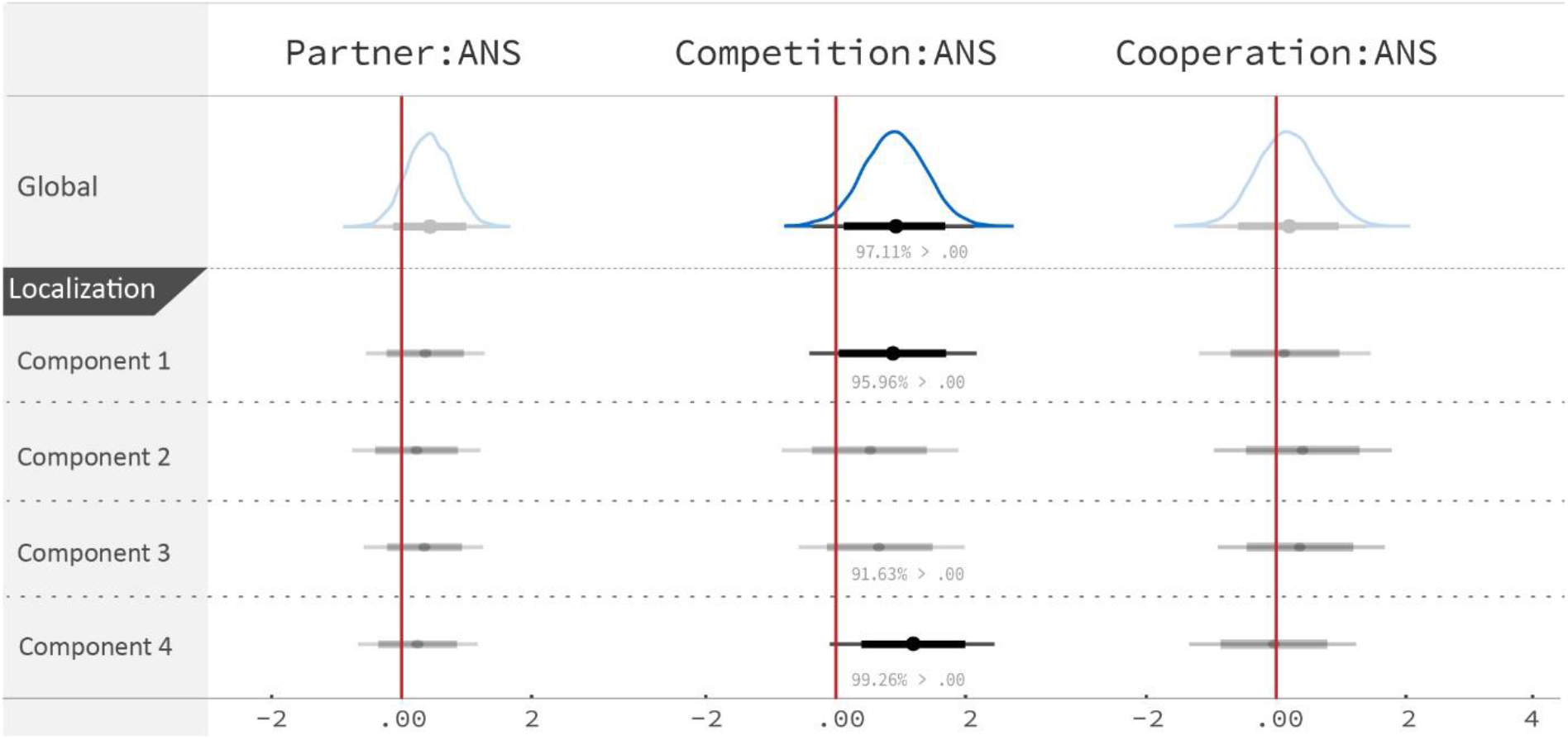
Influences of interpersonal synchrony in the autonomic nervous system (ANS) on neural synchrony (HbO). To investigate whether ANS synchrony predicted increased neural synchrony of mother-child dyads, of competition or cooperation, we examined the interaction effects of ANS synchrony with stranger vs. mother (Partner:ANS), baseline vs. competition (Competition:ANS) and baseline vs. cooperation (Cooperation:ANS). Marginal posterior distributions are plotted for the interaction effects on global and nodal density in the four NMF components. Forest plots show the 99% and 90% two-sided credible intervals (thin and thick black lines) as well as the posterior mean (black dot). Evidence was found for an effect of Competition:ANS on global and nodal density in components 1 and 4. Subsequent analyses showed that only during competition higher density was predicted by higher ANS synchrony. These collective results may indicate that increased interpersonal neural synchrony during competition is related to synchronized arousal, while during cooperation it may go beyond synchrony in ANS signal.

To examine whether this effect was localized, we conducted a multivariate BHM for nodal density. Evidence for an interaction between ANS synchrony and baseline vs. competition was observed in components 1 and 4 as well as some evidence in component 3 (Fig. 5). Again, for baseline vs. cooperation, no interactions were observed with ANS synchrony in any of the components, indicating that increased INS during cooperation was not predicted by increased ANS synchrony. Furthermore, no interactions with partner were found, supporting the notion that increased INS for mother-child compared to stranger-child dyads cannot be attributed to differences in ANS synchrony.

Results for HbR were consistent with the results for HbO, speaking to the validity of the findings (Supplementary Text 2, Supplementary Table 1). Strong and widespread effects of ANS synchrony on global and nodal density were observed for competition, while no effects were found for baseline or cooperation.

In contrast for *behavioral* synchrony, no evidence was found that increased INS was predicted by increased behavioral synchrony (Mean-DRT). Rather for HbO, there was evidence that during competition, less synchronous responses were associated with higher INS (for detailed results see Supplementary Text 8). However, since we were not able to confirm this finding in HbR, this result is not further interpreted.

To summarize, in line with the findings for research question 1, our results showed that increased INS of mother-child dyads was not related to increased behavioral or ANS synchrony. Furthermore, while no relationships between INS, behavioral and ANS synchrony emerged for cooperation, INS and ANS synchrony were positively related during competition.

## 3 Discussion

In this paper we investigated interpersonal synchrony as a multimodal phenomenon ^6,30^ by applying concurrent fNIRS-ECG hyperscanning recordings as well as including behavioral assessments of motor responses in mother-child and stranger-child dyads. First, our results showed increased mother-child relative to stranger-child synchrony only at the neural level. Second, they indicate that increased INS during cooperation cannot be fully explained by ANS and behavioral synchrony, while during competition a positive relationship between INS and ANS synchrony emerged. Together, these results indicate that synchrony occurs across different systems, that the different biobehavioral synchrony markers are clearly differentiable, and that their relationship may be dependent on context.

Our neural findings are generally consistent with those in a sample of 8-18-year-old male children and adolescents, although analytical methods differed ^20^. In this previous study, we found an increased, widespread INS for parent-child competition and more localized effects for parent-child cooperation, however, no increased INS for stranger-child dyads. Here, we examined whether increased INS during competition and cooperation can be attributed to behavioral synchrony, ANS synchrony or other factors. INS was examined at both the global and nodal level.

Based on our results, we are able to rule out a number of possible drivers of the INS that we observed. For example, since INS was higher than in the baseline condition, in which dyads watched a relaxing video together, we rule out the possibility that task-related increases in INS were fully explained by a shared sensory environment. Although we are not able to provide conclusive evidence for a single cause, several possibilities are discussed and evaluated in the light of the present findings.

The first possibility is that aspects of INS may reflect shared social cognitive and attentional processes, including processes of mutual prediction and adaptation ^31^, and the exchange of social ostensive signals ^11,32^. In line thereof, we found that dyads adapted their response times based on feedback more strongly during cooperation than during competition (Supplementary Text 6, Supplementary Table 6). Thus, during cooperation, both adult and child may become entrained to each other as they pay attention to the partner’s behavior, continuously predicting the other’s actions and adapting their own response times based on feedback provided. While during competition, no prediction or adaptation processes are required to successfully complete the task, social comparison processes likely take place during competition, which may potentially lead to an increased INS (see also ^20^).

Specifically, lateral frontopolar cortex regions have been shown to be involved in relational integration, i.e., comparing and integrating information about self and others ^33^. Based on the results of the NMF, showing task-related increases mainly in lateralized nodes for HbO, it can be speculated that comparison processes between one’s own responses and those of the partner facilitate INS. In addition to such cognitive explanations, in animal studies, INS has been found to emerge from two neuronal populations that separately encoded one’s own and the social partner’s behavior ^34^. Thus, both top-down and bottom-up processes may potentially play a role in facilitating INS.

The second possibility is that aspects of INS may arise due to synchronized emotional responses. Here, we found evidence that adult’s and child’s ANS responses become entrained to the task structure (as indicated by the increased autocorrelation every 6 s; Supplementary Text 3, Supplementary Figs. 2 and 3). Further, consistent with previous studies ^16,17^, a significant arousal (ANS) synchrony was observed during both tasks, indicating that cooperation and competition elicit emotional responses in adolescence. However, while no associations between ANS and INS synchrony were found for cooperation, both were positively related during competition, indicating that their association may be dependent on context. In line with our findings for cooperation, a recent study found no relationship between INS and ANS synchrony, measured by respiratory sinus arrhythmia, between 4-6-month-old infants and their mothers when they played with each other ^35^. For competition, the observed relationship my either be explained by the ‘true’ relationship between neural and ANS responses (e.g., ^2,3^) or by false positives, i.e., influences of the ANS on *non-neural* hemodynamic changes (e.g., ^26,27^). Speaking against the latter are the arguments that i) the relationship was only observed during competition and not during cooperation, ii) it was present both in the HbR and HbO signals even though HbR has been found to be less affected by the systemic physiology and cardiac oscillations are less prominent in the HbR signal (e.g., ^27,36^), and iii) the fNIRS connectivity estimator did not include the frequency band of the heart rate.

A third possibility is that INS arises as a result of factors unrelated either to shared cognitive and attentional processes, or to shared ANS entrainment to the task structure. The most consistent aspect of our results was the partner effect (mother-child INS > stranger-child INS). This was observed across conditions (baseline, cooperation, competition), but not found for behavioral and / or ANS synchrony. Instead, shared experiences ^37,38^, social affiliation ^39^, as well as genetic influences ^40^ may lead to higher similarity in brain signals with the mother compared to a stranger. However, since significant heritability is also found for cardiac activities ^41^ and motor reaction times ^42^, the latter explanation seems less likely. Further, this finding is consistent with an increasing number of studies showing higher INS in close relationships, including parent-child days and romantic partners ^13,15,20,43^, and studies indicating that INS may be related to affiliative bonding ^39^. In line thereof, increased similarities in the resting state network connectome of parent-adolescent child dyads have been related to the dyad’s day-to-day emotional synchrony ^38^.

Our study demonstrates the feasibility and utility of multimodal hyperscanning to provide a more holistic view on the neurobiological underpinnings of social interactions. However, one limitation of the study is that with the current fNIRS set-up we focused on the prefrontal cortex, which is important both for social-cognitive and for emotional processing ^44–46^. With recent improvements in fNIRS hardware, future studies may extend the measurements to cover most of the cerebral cortex. This should also be kept in mind when talking about ‘global’ density, which should not be misinterpreted as an effect across the whole brain. Furthermore, with more recent technological developments it is possible to deduct cardiovascular influences originating from the superficial layers of the head (e.g., the skin) from the fNIRS signal by including short distance measurement channels ^47^.

In conclusion, in the current study we introduced multimodal hyperscanning as a novel methodological approach to validate findings and gain a better understanding of the sources of INS. While our results provide support for models which view interpersonal synchrony as a multimodal phenomenon, they also show that synchrony in different behavioral and biological systems should not be considered as a single common factor or unified construct. Instead, results suggest that synchrony in different systems does not necessarily co-occur; rather, that it may reflect distinct processes and that their functional meaning is likely dependent on context. Importantly, we found that increased INS was observed in mother-child compared with stranger-child dyads across conditions (including baseline), and appeared unrelated to increased ANS or behavioral synchrony, suggesting that INS may arise due to neural processes related to social affiliation, going beyond shared arousal and similarities in behavior.

## 4 Methods

### 4.1 Participants

The initial sample consisted of 41 female children, aged between 10 and 18 years, who participated in the study with their biological mothers (mother-child dyads). In addition, each child performed identical tasks with a previously unacquainted female adult (stranger-child dyads). Because INS has been shown to be influenced by the participant’s gender ^9,10^, the current study focused on female children and female adults only. Participants were recruited via previous studies, postings in the intranet of the University Hospital RWTH Aachen as well as flyers. None of the participants had any severe cardiac, neurological or psychiatric conditions.

From the initial sample, one child was excluded because of an attention deficit disorder, two children were excluded due to insufficient fNIRS data quality, one child because of a heart condition and three children because of missing ECG data due to technical errors or insufficient ECG data quality. Thus, the final sample consisted of 34 children (*M* age = 14.26 years, *SD* = 2.206 years, range: 10 – 18 years) and 34 mothers (*M* age = 45.32 years, *SD* = 4.953 years, range: 37 – 56 years). Moreover, a total of 29 female adults served as strangers in the study (*M* age = 23.07 years, *SD* = 2.086 years, range: 19 – 29 years). Of these, 26 adults participated once, one adult twice and two adults three times. Strangers were significantly younger than mothers (*t* (61) = −22.532, *p* < 0.001). For some participants, ECG / fNIRS data were missing in specific experimental conditions mainly due to insufficient data quality or technical errors. Thus, samples sizes varied between *N* = 31 and *N* = 34 for the experimental conditions and measures (for more information see Supplementary Text 9, Supplementary Table 13).

Participants were reimbursed for study participation. The study was approved by the Ethics Committee of the Medical Faculty, University Hospital RWTH Aachen (EK 151/18). All adults, including children of legal age, gave written informed consent for their own study participation as well as, in the case of mothers, for the participation of their child. Children below the age of 18 gave written informed assent.

### 4.2 Procedures

Each experiment began with the baseline condition, followed by the cooperative and competitive tasks. During the experiment, participants were seated next to each other, facing a single computer screen. They were instructed to rest their heads still on a chin rest, in order to reduce movement artifacts, and to refrain from talking to each other. To reduce the participants’ ability to perceive each other’s movements, a towel was placed over their hands. For more details on the procedures, see Supplementary Text 1 and for a video showing the set-up and fNIRS data collection, see ^48^.

A total of 17 children (50%) first completed the three measurements (baseline, cooperation, competition) with the mother and after a short break with the stranger. For 17 children it was the other way around. The order of the cooperative and competitive task was kept constant for both dyads each child was part of but was balanced across children. A total of 7 children (20.6 %) started with mother-child cooperation, 10 children (29.4%) started with mother-child competition, 9 children (26.5%) with stranger-child cooperation and 8 children (23.5%) with stranger-child competition.

### 4.3 Experimental tasks

#### 4.3.1 Baseline

For the baseline condition, a three-minute excerpt from a relaxing aquatic video (Coral Sea Dreaming, Small World Music Inc.) was presented. The aquatic video has been effectively used in previous studies with children to obtain baseline ANS measurements ^49,50^ and served as a low-level control condition to account for the possibility that observed synchronous hemodynamic and physiological changes were due to shared sensory input.

#### 4.3.2 Cooperation and competition task

Adapted versions of the cooperative and competitive computer game tasks of Cui et al. ^19^ were implemented, which have been found appropriate for children ^15,20^. Each player manipulated the on-screen movement of a dolphin towards a ball by pressing a computer key with the goal to either catch the ball together (cooperation) or win the ball for themselves (competition). Each task was composed of two task blocks with 20 trials each and three 30 s rest blocks in alternating order: rest1, task1, rest2, task2, rest3. In line with previous publications ^15,20^, only the two task blocks were considered in the analyses. The trial organization is described in more detail in Supplementary Text 1 and depicted in Supplementary Fig. 1.

During *cooperation*, the goal was to “catch the ball together” by reacting as simultaneously as possible. If the difference in response times was below a predefined threshold, both dolphins jumped to the ball (feedback screen), caught the ball (result screen) and earned a point. If the difference between the response times was above the threshold, only the faster dolphin jumped towards the ball (feedback screen), none of the dolphins caught the ball (result screen) and both participants lost a point. Based on the feedback screen, showing which participant reacted faster and which slower, participants were able to adjust their response times. The temporal threshold was individually adjusted to the response times of the dyad (set to T = 1/8 [RT1+RT2], where RT1 and RT2 indicate the response times of the two participants).

During *competition*, the goal was to “catch and win the ball by oneself” by pressing the response key faster than the other partner. Only the faster dolphin jumped to the ball (feedback screen), caught the ball (result screen) and earned a point while the slower participant lost a point. If the difference between the response times was below 50 ms, both dolphins jumped to the ball (feedback screen), caught the ball (result screen) and gained a point.

### 4.4 Multimodal data acquisition

#### 4.4.1 ECG data acquisition

Electrocardiography (ECG) data were acquired with the Vrije Universiteit Ambulatory Monitoring System (VU-AMS; Netherlands) at a sampling rate of 1000 Hz. In addition, impedance cardiography data were acquired, which is not reported here since it is beyond the scope of the paper. After cleaning the skin with disinfection solution, H98SG, ECG Micropore electrodes (Covidien, Germany) were attached to the participant’s upper body: one slightly below the right collar bone, one on the right side between the lower two ribs and one approximately at the apex of the heart. Prior to the experiment, the internal clock of the VU-AMS device was synchronized to the clock of the stimulation computer, to ensure a temporal synchronization of ECG and fNIRS devices.

#### 4.4.2 fNIRS data acquisition

Functional near-infrared spectroscopy (fNIRS) data were acquired in both subjects simultaneously using a single fNIRS device with a sampling rate of 10 Hz (ETG-4000, Hitachi Medical Corporation, Japan). A ‘3×5’ probe holder grid was mounted to a modified EEG cap (Easycap GmbH, Germany) and probes were inserted into the appropriate holder sockets on the grids. In each grid, eight emitters and seven detectors were positioned alternatingly in three rows, resulting in 22 measurement channels. The source-detector distance was fixed at 3 cm. The caps were placed symmetrically over the participants’ foreheads so that the middle optode of the lowest probe row was placed on the Fpz point of the 10-20 system, and the middle probe column aligned along the sagittal reference curve. The most probable spatial locations of the channels were estimated by the virtual registration method ^51,52^, using the Talairach Daemon ^53^. The brain regions covered by this optode set-up include Brodmann Areas (BAs) 8, 9, 10 and 46 (for the most likely MNI coordinates of the optodes and channels please see: http://www.jichi.ac.jp/brainlab/virtual_registration/Result3x5_E.html). Our recordings concentrated on the prefrontal cortex, since these regions have been frequently found to show significant INS (e.g., ^13,14,34^), and to keep the set-up comparable to previous studies with the same experimental tasks ^15,20^.

### 4.5 Behavioral data analysis

The response times of both participants were recorded during the cooperative and competitive task. As a measure of behavioral synchrony, the mean of the dyad’s absolute differences in their response times (Mean-DRT) was calculated, with smaller values indicating higher synchrony. To further quantify the participants’ task behavior, the number of joint wins during cooperation as well as the number of child’s wins and joint wins during competition are reported in Supplementary Text 5 and Supplementary Table 5. In addition, as an index of how strongly the dyad adapted their response times, we calculated the difference between the mean-DRT of the present trial and its subsequent trial, whereby larger values indicate a stronger adaptation of the dyad. This was calculated for all trials in which participants received feedback showing who responded more quickly (for cooperation this was the case only in trials in which participants had lost a point; Supplementary Text 6, Supplementary Table 6).

### 4.6 ECG data analysis

For the ANS synchrony analyses, we adopted previously used methods ^28,29^ (for further information see Supplementary Text 10). First, R peaks were detected in the raw ECG signal using an automated algorithm. If necessary, R peaks were manually corrected and artifacts removed. Afterwards, for each condition, the IBI time series were resampled at 10 Hz, and the samples were divided into epochs of 2000 ms with fixed on- and offsets to enable an accurate temporal synchronization of the adult’s and child’s IBI values ^54^. An epoch length of 2000 ms was chosen based on minimal amount of time needed to reliably estimate the heart rate ^22^. For each epoch, the mean IBI was computed, resulting in a time series of epoch means for each participant. Artefact removal in the initial IBI series resulted in missing values. Missing values were interpolated with a cubic spline interpolation. If more than 5% of the values were missing of either adult or child in one recording, the respective experimental condition of the dyad was excluded from further analysis (Supplementary Text 9, Supplementary Table 13).

A second order polynomial regression was computed for each epoch means time series in order to remove linear and quadratic trends from the data ^29^. Partial autocorrelation functions (PACFs) showed that the signals had a strong autocorrelation at lag = 1 (Supplementary Text 3, Supplementary Figs. 2 and 3). It is important to remove this autocorrelation, because otherwise spurious correlations may be detected in two independent but autocorrelated time series ^55^. In order to do this, and following an approach used previously ^28,29^, the residuals of the polynomial regression were subjected to Autoregressive Integrated Moving Average (ARIMA) modeling, with one autoregressive term, one moving average term, and integrated noise, and the residuals from this analysis were entered into the cross-correlation calculations. Finally, the cross-correlation at lag = 0 was calculated between the two time series of residuals after ARIMA modeling. Cross-correlations were computed for each condition, and, in case of the cooperation / competition task, for each of the two task blocks. These served as our primary outcome value for ANS synchrony.

In addition, and in order to ensure that the validity of our findings was not specific to the exact measure used to calculate synchrony, we also calculated ANS synchrony by calculating the wavelet coherence, using a method that was as far as possible identical to the method used for calculating INS (Supplementary Text 4).

Of note, both measures used to calculate ANS synchrony do *not* measure how far individual heart beats occur at the same time across the dyad. Rather, they measure how changes in heart rate between consecutive 2000 ms epochs, co-fluctuate across the dyad.

### 4.7 fNIRS data analysis

#### 4.7.1 fNIRS data preprocessing

fNIRS signals were preprocessed by first converting the raw intensity data to optical density data. Second, motion artifacts were detected and reduced by a cubic spline interpolation ^56^. Third, optical density was converted to HbO and HbR concentration changes. The differential pathlength factor was estimated based on the wavelength and the participant’s individual age ^57^. Finally, the data were detrended. Noisy channels were identified based on a semi-automated procedure using several objective criteria in combination with visual inspection and excluded from all subsequent analysis (as described in ^20^). If more than 25% of the channels of a participant in a specific experimental condition was identified as noisy, the complete fNIRS recording was excluded, resulting in missing values (Supplementary Text 9, Supplementary Table 13). For further information on fNIRS data preprocessing see Supplementary Text 11 and Supplementary Table 14.

#### 4.7.2 Connectivity estimator

After signal preprocessing, the statistical dependencies between the dyad’s fNIRS signals were quantified via the bivariate wavelet coherence (WCO). The WCO is a widely applied non-directional functional connectivity estimator, which localizes the signals’ dependencies in the time-frequency space ^58^ and is thereby able to distinguish neural signal components from ANS related frequencies, such as the heart rate. For each signal pair, i.e., for each dyad in each condition and channel combination, the WCO yields a two-dimensional time – ‘frequency’ matrix. These coefficients were then aggregated to a single value, representing the connectivity estimator. To increase the robustness of the estimator, we only considered salient WCO coefficients that are higher than a cut-off value, since these are less affected by noise ^15^. Specifically, we calculated the percentage of salient values across each task block and within a task-related frequency band between 0.08 Hz to 0.5 Hz (period length: 2.02 s - 12.80 s). The task-related frequency band was chosen based on previous studies ^15,20^. It includes the trial duration (~ 7 s for cooperation, ~ 6 s for competition) and importantly lies outside the frequency band of the heart rate (3 Hz – 1 Hz, period length 0.33 s – 1 s). For further information on the WCO and the cut-off calculations see Supplementary Text 12.

#### 4.7.3 Bipartite graph analysis

The complete bipartite graph, *G* = (*V*_1_ ∪ *V*_2_, *E*), was constructed, whereby the fNIRS channels of participant 1, *V*_1_, and of participant 2, *V*_2_, represent the nodes. These two disjoint sets of nodes are connected by edges, *E* ⊆ *V*_1_ × *V*_2_, whose weights *W* are defined by the connectivity estimator (section 4.7.2). Consequently, in hyperscanning the edges can be interpreted as the interpersonal links between the brain regions *V*_1_ of one participant and the brain regions *V*_2_ of another participant. Edges connecting a ‘noisy’ channel were excluded.

In network analysis, it is common practice to exclude edges in order to reduce spurious links and to ensure a more robust network topology. To determine these thresholds, the wavelet coherence was calculated for all possible combinations of independent mother/stranger - child dyads, termed ‘shuffled pairs’, assuming exchangeability of the participant ID while holding the condition and channel combination fixed. Using this blockwise permutation, a shuffled-pair distribution was derived individually for each condition and channel-combination, and the threshold was set to its 95% quantile (for more information see Supplementary Text 13). Thus, only edges were considered which were related to the ‘true’ interaction of the dyad rather than related to random or systemic similarities between brain signals due to the same experimental condition. Based on these reduced graphs both global and nodal graph metrics were calculated.

**Global (inter-brain) density** is defined as the total number of edges, i.e., the interbrain links that survived permutation, relative to the maximum number of possible edges, after noisy channels were excluded ^25^. **Nodal (inter-brain) density** is the number of survived edges for each node that survived permutation, again, relative to the total number of possible edges for the respective node. Thus, nodal density estimates how strongly the temporal activation patterns of a given node are coherent to the temporal activation patterns of the other partner. Thereby it allows to determine the individual contributions of brain regions to this overall connectivity.

#### 4.7.4 NMF

To reduce the dimensionality of the nodal metrics, we applied a NMF. The nodal densities are represented in a matrix 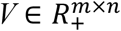, with *m* = 44 nodes, 22 of child and adult, and *n* observations for each dyad in each condition. This matrix was then approximately factorized into a basis matrix 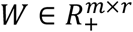 and a coefficient matrix 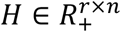, whereby the rank, *r*, is chosen to be smaller than *n* or *m* ^59^. Thus, the matrix H encodes the nodal densities as features for each component, which are later used in the result analysis. The basis matrix W provides the assignment of nodes to components, thus, can be used to look up which combination of nodes contributes to a component-specific effect. To obtain stable results, we performed each NMF with 10,000 iterations. For both HbO and HbR, the rank was chosen to be four, based on the reconstruction error of the original data matrix compared to a shuffled data matrix (for further information see Supplementary Text 14). The nodes which contribute to the four components in term of their weights are depicted in Fig. 3 (HbO) and Supplementary Fig. 4 (HbR).

### 4.8 Validation by shuffled pair analysis

To account for similarities in the dyad’s IBI and fNIRS signals as well as behavioral responses not related to the social interaction, we examined whether synchrony of the actual dyads was higher than synchrony of independent participants involved in the same experimental condition (‘shuffled pairs’, see section 4.7.3). To this end, interpersonal synchrony measures were calculated for all possible shuffled mother /stranger - child pairs.

Specifically, sets of shuffled pairs were constructed for each child by varying the adult partner and for each adult by varying the child partner, while holding the condition fixed. Next, a dyad- and condition- specific mean shuffled pair synchrony value was derived by averaging across the synchrony values of the child’s and adult’s shuffled pair sets in each condition. Thus, for each dyad, we obtained one actual synchrony value and one mean shuffled pair synchrony value ^20^.

While shuffled pairs performed the same cooperative / competitive task, the timing of the trials and length of task blocks differed between subjects due to the variable inter-trial interval and the subjects’ responses. Since ANS and neural synchrony analysis requires an equal length of the signals, the longer task block was cut at the end to have the same length as the shorter task block.

### 4.9 Bayesian result analysis

To derive an estimate of how the experimental conditions affect interpersonal synchrony in different systems and the factors influencing INS, BHMs were used. This method is gaining increasing importance for neuroscience and brain network analyses ^60^. The Bayesian framework comes with several advantages compared to the classical frequentist approach. Bayesian models allows to incorporate prior knowledge about the parameters in the models and to specify different response distributions. This is particularly important since nodal density follows a non-Gaussian distribution with long tails towards high values, thereby violating assumptions of many classical frequentist tests ^61^. Furthermore, the BHM does not rely on *p*-values but derives a probability statement for each of the parameters of interest. In the result section, we report the mean of the parameter’s estimated marginal posterior distribution as well as its two-sided 90% CI, which is defined as the probabilistic interval that is believed to contain a given parameter ^62^. For discussion purposes, two-sided 90% CIs which do not include zero are interpreted as statistical evidence for a given effect, two-sided 80% CIs are interpreted as weak evidence for an effect. Prior to the BHM analyses, missing fNIRS and ANS synchrony values were imputed by multiple imputation. For more information on the imputations, BHM implementations and quality checks see Supplementary Text 15.

#### 4.9.1 In which systems does interpersonal synchrony occur?

First, to examine whether actual pairs differed from shuffled pairs, we calculated individual BHMs for i) INS, ii) ANS and iii) behavioral synchrony with global density, ANS cross-correlations and Mean-DRT as the response variable, respectively. The models included pair (0 = shuffled, 1 = actual), experimental condition as well as their interaction as predictors. Reported are the effects of pair for each experimental condition.

Second, to directly compare the different experimental conditions, BHMs were again calculated for i) INS, ii) ANS and iii) behavioral synchrony. For models i) and ii), predictors included: competition (0 = baseline, 1 = competition), cooperation (0 = baseline, 1 = cooperation), partner (0 = stranger, 1 = mother), as well as the two-way interactions between competition / cooperation and partner. For model iii), task was coded with 0 = cooperation and 1 = competition. If there is no evidence for an interactive effect, unconditional main effects are reported.

For nodal density, we calculated multivariate BHMs, which included all four NMF components as response variables. It should be noted that BHMs integrate all effects into one model and thereby address multiple comparison issues ^63^.

#### 4.9.2 Which factors predict interpersonal neural synchrony?

Second, we examined whether ANS and behavioral synchrony predicted INS. To this end, we calculated univariate or multivariate BHMs for global and nodal density, respectively. First, we estimated the effects of ANS synchrony, competition (0 = baseline, 1 = competition), cooperation (0 = baseline, 1 = competition), partner (0 = stranger, 1 = mother), as well as their two-way interactions with ANS synchrony. To calculate cross-level interactions, in this case with ANS synchrony (level 1) nested in task and partner (level 2), it is advisable to conduct a group-mean centering of the level 1 predictor prior to the analysis ^64^. Thus, ANS synchrony values were group-mean centered by subtracting the mean value in the respective experimental condition. Equivalent (multivariate) BHMs were formulated for behavioral synchrony, estimating the effects of task (0 = competition, 1 = cooperation), partner (0 = stranger, 1 = mother), behavioral synchrony (group-mean centered) and their two-way interactions on global and nodal density.

## Acknowledgments

The authors would like to thank the participants and are grateful to Yasemin Capraz for her help with data collection. We thank Helena Oldenhof and Giorgia Banzato for their advice on ECG data collection and preprocessing.

## Author contributions

V.R. and K.K. designed the study and acquired funding. V.R., S.Wi., C.L.W. collected and preprocessed the data. V.R., C.G., K.K., S.Wa., V.L. and W.S. contributed to the data analysis and interpretation. V.R. and C.G. performed the computations. V.R., C.G., S.Wa. drafted the paper. All authors have revised the paper.

## Competing Interests

The authors declare no competing financial interests.

## Funding

This work was funded by the Excellence Initiative of the German federal state and governments (ERS Seed Fund, OPSF449) and the START-Programme of the medical faculty of the RWTH Aachen University. The Hitachi NIRS system was supported by a funding of the German Research Foundation DFG (INST 948/18-1 FUGG). Scientific computations were partly performed with the computing resources granted by RWTH Aachen University under project 2082. The work of C.G. was performed as part of the Helmholtz School for Data Science in Life, Earth and Energy (HDS-LEE).

## Notes

### Competing Interest Statement

The authors have declared no competing interest.

## References

1 Damasio, A. Descartes’ error: Emotion, Reason, and the Human Brain. (G.P. Putnam’s Son, New York, 1994.

2 Critchley, H. D. et al. Activity in the human brain predicting differential heart rate responses to emotional facial expressions. NeuroImage 24, 751–762 (2005).

3 Lane, R. D. et al. Neural correlates of heart rate variability during emotion. NeuroImage 44, 213–222 (2009).

4 Barber, A. D. et al. Parasympathetic arousal-related cortical activity is associated with attention during cognitive task performance. NeuroImage 208, 116469 (2020).

5 Breeden, A., Siegle, G., Norr, M., Gordon, E. & Vaidya, C. Coupling between spontaneous pupillary fluctuations and brain activity relates to inattentiveness. Eur. J. Neurosci. 45, 260–266 (2017).

6 Hari, R., Henriksson, L., Malinen, S. & Parkkonen, L. Centrality of social interaction in human brain function. Neuron 88, 181–193 (2015).

7 Hasson, U., Ghazanfar, A. A., Galantucci, B., Garrod, S. & Keysers, C. Brain-to-brain coupling: a mechanism for creating and sharing a social world. Trends Cogn. Sci. 16, 114–121 (2012).

8 Davis, M., West, K., Bilms, J., Morelen, D. & Suveg, C. A systematic review of parent– child synchrony: it is more than skin deep. Dev. Psychobiol. 60, 674–691 (2018).

9 Baker, J. M. et al. Sex differences in neural and behavioral signatures of cooperation revealed by fNIRS hyperscanning. Sci. Rep. 6, 26492 (2016).

10 Cheng, X., Li, X. & Hu, Y. Synchronous brain activity during cooperative exchange depends on gender of partner: a fNIRS-based hyperscanning study. Hum. Brain Mapp. 36, 2039–2048 (2015).

11 Leong, V. et al. Speaker gaze increases information coupling between infant and adult brains. Proc. Natl. Acad. Sci. 114, 13290–13295 (2017).

12 Nguyen, T. et al. The effects of interaction quality on neural synchrony during mother-child problem solving. Cortex 124, 235–249 (2020).

13 Pan, Y., Cheng, X., Zhang, Z., Li, X. & Hu, Y. Cooperation in lovers: an fNIRS-based hyperscanning study. Hum. Brain Mapp. 38, 831–841 (2017).

14 Piazza, E. A., Hasenfratz, L., Hasson, U. & Lew-Williams, C. Infant and adult brains are coupled to the dynamics of natural communication. Psychol. Sci. 31, 6–17 (2020).

15 Reindl, V., Gerloff, C., Scharke, W. & Konrad, K. Brain-to-brain synchrony in parent-child dyads and the relationship with emotion regulation revealed by fNIRS-based hyperscanning. NeuroImage 178, 493–502 (2018).

16 Järvelä, S., Kivikangas, J. M., Kätsyri, J. & Ravaja, N. Physiological linkage of dyadic gaming experience. Simul. Gaming 45, 24–40 (2014).

17 Chanel, G., Kivikangas, J. M. & Ravaja, N. Physiological compliance for social gaming analysis: cooperative versus competitive play. Interact. Comput. 24, 306–316 (2012).

18 Balconi, M. & Vanutelli, M. E. Cooperation and competition with hyperscanning methods: review and future application to emotion domain. Front. Comput. Neurosci. 11, 86 (2017).

19 Cui, X., Bryant, D. M. & Reiss, A. L. NIRS-based hyperscanning reveals increased interpersonal coherence in superior frontal cortex during cooperation. NeuroImage 59, 2430–2437 (2012).

20 Kruppa, J. A. et al. Brain and motor synchrony in children and adolescents with ASD - a fNIRS hyperscanning study. Soc. Cogn. Affect. Neurosci. 16, 103 – 116 (2021).

21 Dai, B. et al. Neural mechanisms for selectively tuning in to the target speaker in a naturalistic noisy situation. Nat. Commun. 9, 2405 (2018).

22 Helm, J. L., Miller, J. G., Kahle, S., Troxel, N. R. & Hastings, P. D. On measuring and modeling physiological synchrony in dyads. Multivariate Behav. Res. 53, 521–543 (2018).

23 Gerloff, C., Konrad, K., Bzdok, D., Büsing, C. & Reindl, V. Interacting brains revisited: a cross-brain network neuroscience perspective. Preprint at: https://biorxiv.org/cgi/content/short/2021.02.20.432051v1 (2021).

24 Ciaramidaro, A. et al. Multiple-brain connectivity during third party punishment: an EEG hyperscanning study. Sci. Rep. 8, 6822 (2018).

25 Santamaria, L. et al. Emotional valence modulates the topology of the parent-infant inter-brain network. NeuroImage 207, 116341 (2020).

26 Chen, J. E. et al. Resting-state “physiological networks”. NeuroImage 213, 116707 (2020).

27 Tachtsidis, I. & Scholkmann, F. False positives and false negatives in functional near-infrared spectroscopy: issues, challenges, and the way forward. Neurophotonics 3, 031405 (2016).

28 Feldman, R., Magori-Cohen, R., Galili, G., Singer, M. & Louzoun, Y. Mother and infant coordinate heart rhythms through episodes of interaction synchrony. Infant Behav. Dev. 34, 569–577 (2011).

29 Suveg, C., Shaffer, A. & Davis, M. Family stress moderates relations between physiological and behavioral synchrony and child self-regulation in mother–preschooler dyads. Dev. Psychobiol. 58, 83–97 (2016).

30 Semin, G. R. in Social psychology: handbook of basic principles (eds A. W. Kruglanski & E. T. Higgins) 630 – 649 (The Guilford Press, 2007).

31 Hamilton, A. F. d. C. Hyperscanning: beyond the hype. Neuron 109, 404 – 407 (2020).

32 Wass, S. V., Whitehorn, M., Marriot Haresign, I., Phillips, E. & Leong, V. Interpersonal neural entrainment during early social interaction. Trends Cogn. Sci. 24, 329 – 342 (2020).

33 Raposo, A., Vicens, L., Clithero, J. A., Dobbins, I. G. & Huettel, S. A. Contributions of frontopolar cortex to judgments about self, others and relations. Soc. Cogn. Affective Neurosci. 6, 260–269 (2011).

34 Kingsbury, L. et al. Correlated neural activity and encoding of behavior across brains of socially interacting animals. Cell 178, 429–446.e416 (2019).

35 Nguyen, T., Abney, D. H., Salamander, D., Bertenthal, B. & Hoehl, S. Social touch is associated with neural but not physiological synchrony in naturalistic mother-infant interactions. Preprint at: https://www.biorxiv.org/content/10.1101/2021.01.21.427664v1 (2021).

36 Kirilina, E. et al. Identifying and quantifying main components of physiological noise in functional near infrared spectroscopy on the prefrontal cortex. Front. Hum. Neurosci. 7, 864 (2013).

37 Gabard-Durnam, L. J. et al. Stimulus-elicited connectivity influences resting-state connectivity years later in human development: A prospective study. J. Neurosci. 36, 4771–4784 (2016).

38 Lee, T.-H., Miernicki, M. E. & Telzer, E. H. Families that fire together smile together: Resting state connectome similarity and daily emotional synchrony in parent-child dyads. NeuroImage 152, 31–37 (2017).

39 Zheng, L. et al. Affiliative bonding between teachers and students through interpersonal synchronisation in brain activity. Soc. Cogn. Affective Neurosci. 15, 97 – 109 (2020).

40 Glahn, D. C. et al. Genetic control over the resting brain. Proc. Natl. Acad. Sci. 107, 1223–1228 (2010).

41 Muñoz, M. L. et al. Heritability and genetic correlations of heart rate variability at rest and during stress in the Oman Family Study. J. Hypertens. 36, 1477 – 1485 (2018).

42 Kuntsi, J. et al. Reaction time, inhibition, working memory and ‘delay aversion’ performance: genetic influences and their interpretation. Psychol. Med. 36, 1613 – 1624 (2006).

43 Kinreich, S., Djalovski, A., Kraus, L., Louzoun, Y. & Feldman, R. Brain-to-brain synchrony during naturalistic social interactions. Sci. Rep. 7, 17060 (2017).

44 Goldin, P. R., McRae, K., Ramel, W. & Gross, J. J. The neural bases of emotion regulation: reappraisal and suppression of negative emotion. Biol. Psychiatry 63, 577–586 (2008).

45 Decety, J., Jackson, P. L., Sommerville, J. A., Chaminade, T. & Meltzoff, A. N. The neural bases of cooperation and competition: an fMRI investigation. NeuroImage 23, 744–751 (2004).

46 Amodio, D. M. & Frith, C. D. Meeting of minds: the medial frontal cortex and social cognition. Nat. Rev. Neurosci. 7, 268–277 (2006).

47 Nozawa, T., Sasaki, Y., Sakaki, K., Yokoyama, R. & Kawashima, R. Interpersonal frontopolar neural synchronization in group communication: an exploration toward fNIRS hyperscanning of natural interactions. NeuroImage 133, 484–497 (2016).

48 Reindl, V. et al. Conducting hyperscanning experiments with functional near-Infrared spectroscopy. J. Vis. Exp. 143, e58807 (2019).

49 Prätzlich, M. et al. Resting autonomic nervous system activity is unrelated to antisocial behaviour dimensions in adolescents: cross-sectional findings from a European multi-centre study. J. Crim. Justice 65, 101536 (2018).

50 Piferi, R. L., Kline, K. A., Younger, J. & Lawler, K. A. An alternative approach for achieving cardiovascular baseline: viewing an aquatic video. Int. J. Psychophysiol. 37, 207–217 (2000).

51 Singh, A. K., Okamoto, M., Dan, H., Jurcak, V. & Dan, I. Spatial registration of multichannel multi-subject fNIRS data to MNI space without MRI. NeuroImage 27, 842–851 (2005).

52 Tsuzuki, D. et al. Virtual spatial registration of stand-alone fNIRS data to MNI space. NeuroImage 34, 1506–1518 (2007).

53 Lancaster, J. L. et al. Automated Talairach atlas labels for functional brain mapping. Hum. Brain Mapp. 10, 120–131 (2000).

54 Berntson, G. G., Cacioppo, J. T. & Quigley, K. S. The metrics of cardiac chronotropism: biometric perspectives. Psychophysiology 32, 162–171 (1995).

55 Dean, R. T. & Dunsmuir, W. T. Dangers and uses of cross-correlation in analyzing time series in perception, performance, movement, and neuroscience: the importance of constructing transfer function autoregressive models. Behav. Res. Methods 48, 783–802 (2016).

56 Scholkmann, F., Spichtig, S., Muehlemann, T. & Wolf, M. How to detect and reduce movement artifacts in near-infrared imaging using moving standard deviation and spline interpolation. Physiol. Meas. 31, 649 – 662 (2010).

57 Scholkmann, F. & Wolf, M. General equation for the differential pathlength factor of the frontal human head depending on wavelength and age. J. Biomed. Opt. 18, 105004 (2013).

58 Daubechies, I. The wavelet transform, time-frequency localization and signal analysis. IEEE Trans. Inf. Theory 36, 961–1005 (1990).

59 Lee, D. D. & Seung, H. S. Algorithms for non-negative matrix factorization In Advances in neural information processing systems. 556–562 (2001).

60 Bzdok, D., Floris, D. L. & Marquand, A. F. Analysing brain networks in population neuroscience: a case for the Bayesian philosophy. Philos. Trans. R. Soc. Lond. B Biol. Sci. 375, 20190661 (2020).

61 Bullmore, E. & Sporns, O. Complex brain networks: graph theoretical analysis of structural and functional systems. Nat. Rev. Neurosci. 10, 186–198 (2009).

62 Aczel, B. et al. Discussion points for Bayesian inference. Nat. Hum. Behav. 4, 561 – 563 (2020).

63 Gelman, A., Hill, J. & Yajima, M. Why we (usually) don’t have to worry about multiple comparisons. J. Res. Educ. Eff. 5, 189–211 (2012).

64 Enders, C. K. & Tofighi, D. Centering predictor variables in cross-sectional multilevel models: a new look at an old issue. Psychol. Methods 12, 121 – 138 (2007).

